# mRNA-LNP vaccines against Hepatitis B virus induce protective immune responses in preventive and chronic mouse challenge models

**DOI:** 10.1101/2025.02.12.636533

**Authors:** María José Limeres, Rocio Gambaro, Malin Svensson, Silvia Fraude-El Ghazi, Leah Pretsch, Daniel Frank, German A. Islan, Ignacio Rivero Berti, Matthias Bros, Ying K. Tam, Hiromi Muramatsu, Norbert Pardi, Stephan Gehring, Maximiliano L. Cacicedo

**Author notes:** Correspondence: Maximiliano L. Cacicedo, Stephan Gehring, Norbert Pardi.

## Abstract

Over 300 million people worldwide suffer from chronic hepatitis B virus infections that can cause serious liver damage and hepatocellular carcinoma. Ineffective innate and adaptive immune responses characterize these chronic infections, making the development of a therapeutic vaccine an urgent medical need. While current vaccines can prevent HBV infections, they are ineffective in treating chronic disease. This study investigated lipid nanoparticle (LNP)-formulated nucleoside-modified mRNA vaccines encoding Hepatitis B surface antigen (HBsAg) for prophylactic and therapeutic applications. We found that HBsAg mRNA-LNP vaccines induced robust humoral and cellular immune responses, outperforming the protein-based vaccine approved for human use. The incorporation of an MHC class I signal peptide further enhanced Th1-biased responses preventing HBV infections in a mouse model. Importantly, mRNA-LNP vaccination led to seroconversion, HBsAg clearance, and strong T cell responses in a chronically infected mouse model. These findings highlight the potential of mRNA-LNP as an alternative and effective vaccine modality for HBV prophylaxis and therapeutic use in treating chronic infections.

## Introduction

Hepatitis B virus (HBV) is a small enveloped DNA virus that selectively infects human hepatocytes.^1^ Upon infection, the circular viral genome (relaxed circular DNA) is transported to the nucleus of the infected cell where host enzymes convert it into an episomal transcriptional template (covalently closed circular DNA, cccDNA) leading to chronic infection.^2,3^ Even though effective prophylactic anti-hepatitis B vaccines exist, there are an estimated 300 million people worldwide who suffer from chronic hepatitis B (CHB) infections, and ∼1.5 million new infections occur each year.^2,3^ The principal route of HBV transmission in high-endemic countries is from mother to child; the most relevant transmission routes in low-endemic countries are sexual and parenteral. Recently, mother-to-child transmission has been reduced significantly by administering a prophylactic vaccine during the first 24 hours of life.^4^ In many cases, however, the induction of protective immunity, assessed in terms of anti-hepatitis B surface antigen (HBsAg) levels of 10-100 IU/L in low-responders and anti-HBsAg levels ≥100 IU/L in high-responders, is insufficient, and breakthrough infections occur in 5–10% of vaccinated individuals. Unvaccinated children infected with HBV, on the other hand, exhibit a 90% probability of developing chronic infections and a 25% chance of eventually suffering liver damage.^5^ Pediatric CHB patients often fail to exhibit symptoms during childhood but carry a high risk of developing hepatocellular carcinoma (HCC) as adults.^6^ In fact, CHB is a major cause of HCC.^1^

CHB infections are characterized by ineffective innate and adaptive immune responses. HBV modulates innate immunity leading to an inability to mount an adaptive immune response in chronic HBV-infected patients, resulting in viral persistence.^7–9^ In summary, CHB infection is characterized by T cell exhaustion with reduced cytotoxic activity, impaired cytokine production, and sustained expression of cell-surface inhibitory receptors.^10^

High levels of serum HBsAg found in most CHB patients impair B and T cell responses. It has been suggested that treatment options that clear systemic HBsAg levels could reverse immune dysfunction.^11^ The concept of “functional cure” is used in this context to describe the loss of detectable serum HBsAg with or without associated seroconversion to anti-HBsAg antibody production.^4^ Unfortunately, functional cures are not achieved with current treatment options that include the use of pegylated IFN-α and nucleoside analogs. In the absence of effective CHB treatments, there is an urgent medical need for novel interventions and immunotherapies that restore life-long patient immunity.

The coordinated activation of anti-HBV-specific humoral and cell-mediated immunity in patients who completely recover from HBV infections indicates that immune activation and functional cure are achievable.^12,13^ B cells capable of secreting anti-HBV antibodies and robust helper and cytotoxic T cell responses observed in patients who recovered indicate that therapeutic approaches that elicit anti-viral immune mechanisms could induce a functional cure.

Prophylactic vaccines tested for their therapeutic effects in CHB patients achieved little success, i.e., the induction of specific antibody production, but no cytotoxic T-cell response.^14,15^ New vaccine formulations showed only limited success when used curatively.^16–18^ A therapeutic vaccine is needed that can induce robust innate and adaptative immunity, optimized by incorporating the appropriate adjuvants into the vaccine formulation.^19^

Lipid nanoparticle (LNP)-encapsulated nucleoside-modified mRNA vaccines are well-tolerated and, in many cases, have proven to be more effective than conventional vaccine platforms.^20^ The capacity to generate both humoral and cell-mediated immunity makes mRNA-LNP-based vaccine strategies a feasible option to address problematic infections such as CHB. Here, two potential anti-hepatitis B vaccine candidates were generated using either the native HBsAg sequence or a second construct with an incorporated MHC class I signal peptide (SP) to enhance the routing of the protein product through MHC class I intracellular compartments. While both vaccines mounted a robust immune response and outperformed the currently used protein-based anti-HBV vaccine, the *SP-HBsAg* mRNA-LNP vaccine was superior in preventing HBV infections in a mouse model by inducing a robust antigen-specific Th1 cell-biased immune response. On the other hand, both vaccines decreased serum hepatitis B surface antigen (HBsAg) in chronically infected mice. Remarkably, CHB mice immunized with a combination of both mRNA vaccine candidates generated the strongest CD8^+^ T cell-mediated response while preserving low HBsAg plasma levels. These findings suggest that nucleoside-modified mRNA-LNP vaccines represent an alternative, more effective anti-hepatitis B vaccine modality for prophylactic and therapeutic use.

## Results

### Human monocyte-derived dendritic cells (hmoDCs) transfected with *HBsAg* mRNA produce HBsAg protein

The presence of intracellular and extracellular HBsAg protein in hmoDCs transfected with *HBsAg* mRNA or incubated with HBsAg protein as control was assessed and compared by flow cytometric analysis. A marked increase in the frequency of intracellular HBsAg-positive hmoDCs occurred in both cases; a significantly higher signal was observed, however, in the *HBsAg* mRNA-transfected cells (Figures 1a and 1b). Moreover, only *HBsAg* mRNA-transfected hmoDCs exhibited a significant signal for HBsAg protein stained extracellularly (Figure 1c and 1d). Furthermore, *HBsAg* mRNA-transfected hmoDCs exhibited an elevated expression of cell surface activation markers, i.e., HLA-DR, CD40, CD80 and CD86, measured by flow cytometry (Figure 1e, Figure S1). Transfection with ovalbumin-encoding mRNA (*ova* mRNA), an irrelevant mRNA control, exhibited no immunostimulatory effect. Similarly, incubation with neither HBsAg protein nor ENGERIX^®^-B, a licensed anti-hepatitis B vaccine, exerted a significant effect on the activation markers expressed by hmoDCs.

**Figure 1.**
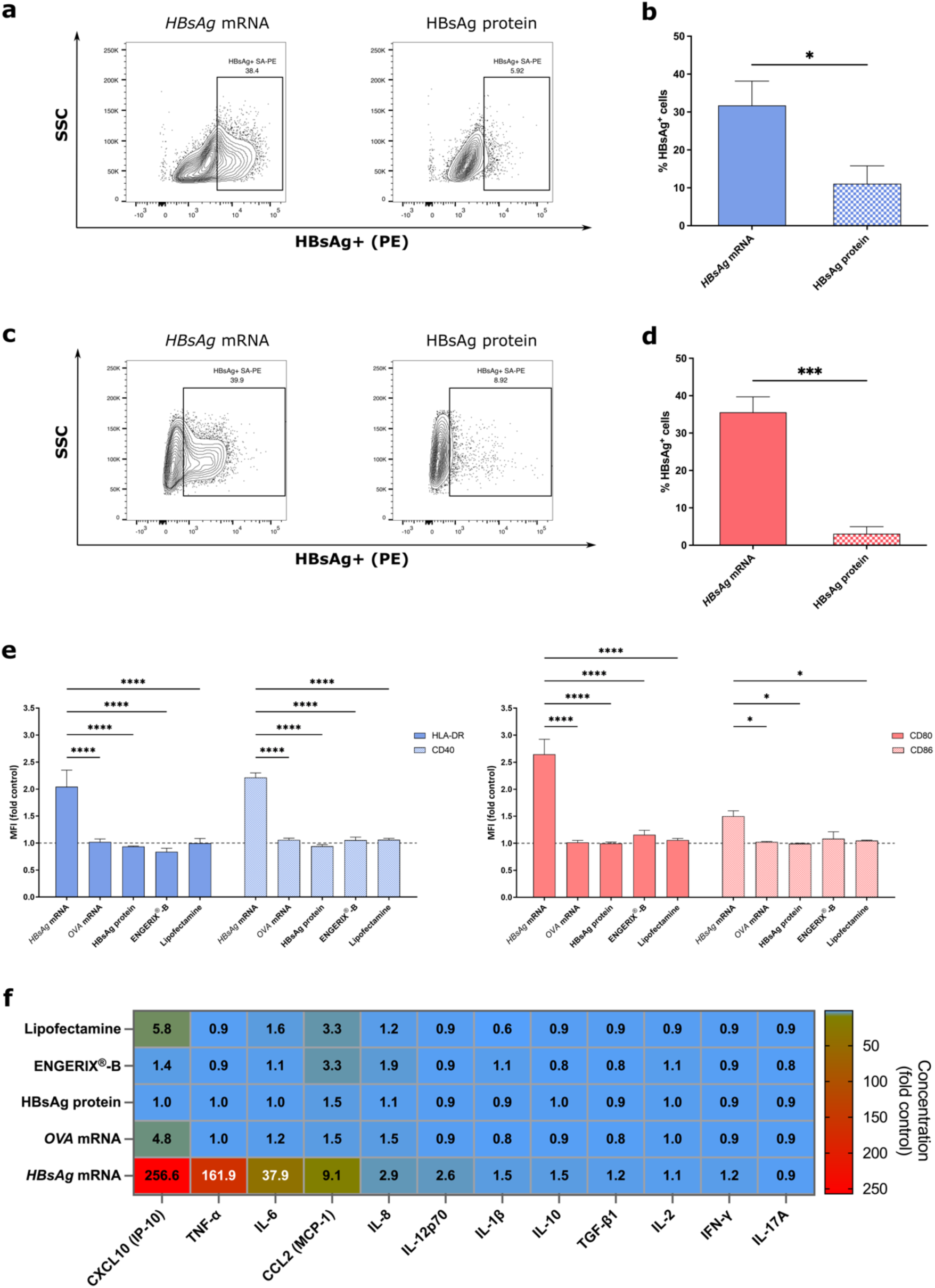
*HBsAg* mRNA stimulates activation and antigen production by hmoDCs *in vitro*. Intracellular (a, b) and extracellular (c, d) HBsAg synthesized by transfected hmoDCs was evaluated by flow cytometry. The hmoDCs incubated with HBsAg protein were used as a control. Background signals from untreated controls were extracted from the depicted data. (e) The expression of HLA-DR, CD40, CD80 and CD86 activation markers by hmoDCs transfected with *HBsAg* mRNA was evaluated by flow cytometry. *Ova* mRNA was used as an irrelevant mRNA control. ENGERIX^®^-B, HBsAg protein and Lipofectamine Messenger Max were also used for control comparisons. The mean fluorescence intensity (MFI) of treated and untreated control groups was compared. Results are given as the means ± SEM obtained in 4 independent experiments. Significantly different compared to the control group: *p <0.05; ***p <0.001, ****p <0.0001 (two-tailed Student’s *t*-test and two-way ANOVA, Tukey’s multiple comparisons test). (f) Cell culture supernates were collected for cytokine contents. Heatmap represents the fold increase in cytokine/chemokine concentrations in supernates obtained from treated samples, compared to the untreated control.

Cytokines and chemokines secreted into the cell culture supernates were also quantified (Figure 1f). The hmoDCs transfected with *HBsAg* mRNA generated large amounts of CXCL10 (IP-10) and TNF-α compared to non-transfected hmoDCs, the concentrations of IL-6 and CCL2 (MCP-1) were slightly elevated; none of the other cytokines or chemokines included in the panel was detectable. *Ova* mRNA transfection, or incubation with either HBsAg protein or ENGERIX^®^-B, failed to stimulate the secretion of any cytokines or chemokines by hmoDCs.

### An *HBsAg* mRNA-LNP vaccine promotes antigen-specific antibody production and a shift towards a Th-1 type response

Mice were injected intramuscularly (i.m.) with 5 μg *HBsAg* mRNA-LNP and early immune-related effects were evaluated 24 hours after administration (Figure S2a-b). The activation of T cells and DCs isolated from spleens and inguinal lymph nodes (iLNs) was evaluated. Expression of the early activation marker, CD69, by CD4^+^ and CD8^+^ T cells was upregulated significantly in mice vaccinated with *HBsAg* mRNA-LNP (Figure S2c). Similarly, the DCs in the spleen and draining lymph nodes of vaccinated animals exhibited marked increases in CD80 and CD86 expression (Figure S2d).

The endoplasmic reticulum (ER), early and late endosomes, and lysosomes have been described as part of the essential machinery for antigen processing and presentation by MHC class I and class II molecules.^21^ Different studies have shown that engineering the antigenic sequence by attaching endoplasmic reticulum translocation signal sequences to the N-terminus can improve antigen presentation and, therefore, optimize specific immune responses.^22–24^ Furthermore, introducing modifications into the C-terminus of the antigen has also been studied in an effort to improve antigen trafficking. Signal motifs such as lysosome-associated membrane proteins (LAMPs) and the MHC class I trafficking domain (MITD) linked to C-terminal regions of the antigen, combined with N-terminal signal peptides, have been reported to strongly enhance CTL responses.^24–28^ Therefore, to optimize the generation of HBsAg-specific immunity by mRNA-LNP, trafficking sequences were incorporated into the *HBsAg* mRNA construct to enhance antigen presentation by optimizing routing through MHC class I intracellular compartments (Figure 2a). Specifically, an MHC class I signal peptide (SP) was included in the 5’ region of the mRNA construct, upstream of the HBsAg sequence. Additionally, the sequence coding for MITD was incorporated into the 3’ region downstream of the HBsAg sequence. A construct with the 5’ SP sequence, but without 3’ MITD was also generated and tested.

**Figure 2.**
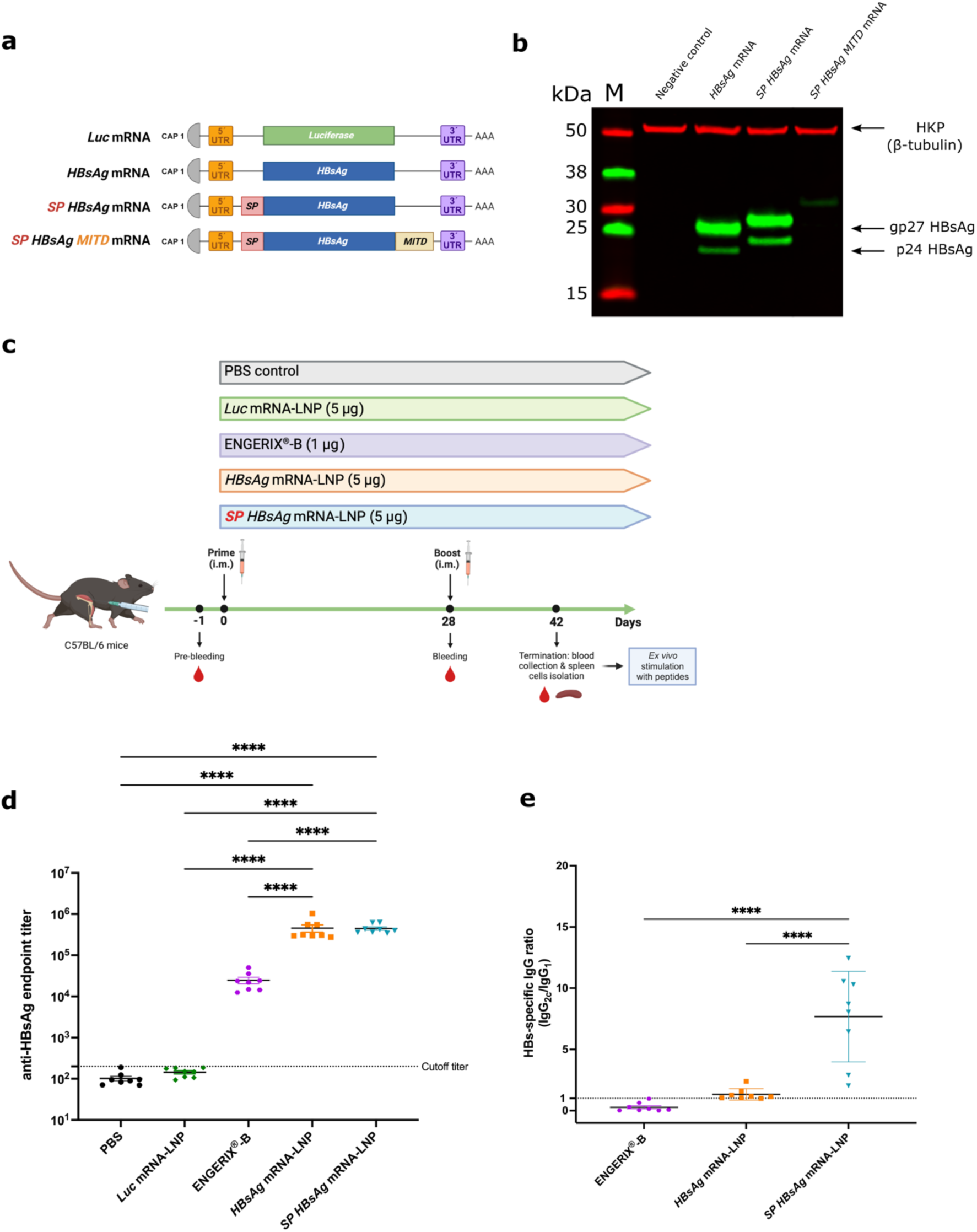
Antigen optimized by incorporating signal peptides into mRNA constructs results in higher anti-HBsAg antibody titers. (a) Three *HBsAg* nucleoside-modified mRNA-LNP vaccines were prepared by modifying the antigenic sequence: (i) a wild-type HBsAg construct (*HBsAg* mRNA); (ii) the *HBsAg* mRNA incorporating an MHC class I signal peptide (SP) included in the 5’ region (*SP-HBsAg* mRNA); and (iii) a construct with the 5’ SP and a transmembrane and cytosolic trafficking domain of MHC class I (MITD) added to the 3’ region of the *HBsAg* sequence (*SP-HBsAg-MITD* mRNA). *Firefly luciferase* mRNA (*Luc* mRNA) was prepared and used as an irrelevant mRNA control. (b) Western blot analysis of HepG2 cell lysates after transfection with mRNA constructs. Non-transfected HepG2 cells served as a negative control. HBsAg protein is expressed as glycosylated (gp27) and non-glycosylated (p24) isomers. (c) Experimental design. Mice were vaccinated with mRNA-LNP constructs on day 0 (prime). Four weeks after priming, the second dose (boost) was injected (d28). Vaccination with ENGERIX^®^-B was used as a positive control, *Luc* mRNA-LNP and PBS were used as negative controls. The experiment was terminated on day 42, and blood and spleen were collected for analyses. Blood was also collected on day -1 (pre-immune bleed) and day 28 prior to boosting. (d) Total anti-HBsAg IgG titers at termination (day 42). (e) IgG2c/IgG1 ratio at termination. Data are given as the means ± SEM for animal cohorts (each 8 mice/group). Statistical differences: ****p <0.0001 (one-way ANOVA, Tukey’s multiple comparisons test).

The translation efficiency of the different mRNA constructs was evaluated by Western blot analysis using the protein extract from mRNA-transfected HepG2 cells. While *HBsAg* and *SP-HBsAg* mRNA-transfected cells produced a significant amount of protein, the presence of MITD in the 3’ region of the construct dramatically decreased protein synthesis (Figure 2b). Predictably, the incorporation of SP resulted in the generation of a protein that was slightly larger in size. The construct containing the MITD modification was not evaluated in immunization experiments due to its poor translation efficiency.

Experiments were undertaken to characterize the HBsAg-specific humoral responses of C57BL/6Ncr mice vaccinated twice (days 0 and 28) i.m. with 5 μg mRNA-LNP vaccines (Figure 2c). *SP-HBsAg* and *HBsAg* mRNA-LNP vaccines were evaluated in the immunization studies. Mice immunized with *Luc* mRNA-LNP, ENGERIX^®^-B (1 μg) and PBS were used as controls. ENGERIX^®^-B dose (1 μg) was selected based on a 20x reduction of the recommended human dose (20 μg).^29^ The same ratio was applied for mRNA-LNP vaccines (5 μg) compared to the Moderna COVID-19 (mRNA-1273) vaccine that was initially 100 μg mRNA.^30^ Two weeks after vaccine boost, HBsAg-specific antibody titers were quantified. While ENGERIX^®^-B vaccination resulted in modest antibody levels, mice vaccinated with either *HBsAg* mRNA-LNP or *SP-HBsAg* mRNA-LNP exhibited marked increases in antigen-specific endpoint IgG levels (Figure 2d). The ratio of IgG2c to IgG1 subtypes was determined to assess the balance between Th1- and Th2-biased immune responses. Only *SP-HBsAg* mRNA-LNP induced a shift towards a Th1-type response demonstrated by an average IgG2c/IgG1 ratio significantly greater than 1 (Figure 2e).

T cell responses were further characterized by intracellular cytokine staining (ICS) and analyzed by flow cytometry. Unlike the positive control licensed vaccine, mRNA-LNP vaccines elicited HBsAg-specific CD4^+^ and CD8^+^ T cells that expressed IFN-γ, TNF-α and IL-2 (Figures 3a and 3b, Figure S3). In the cases of both CD4^+^ and CD8^+^ T cells, mRNA-LNP outperformed the responses generated by vaccination with ENGERIX^®^-B.

**Figure 3.**
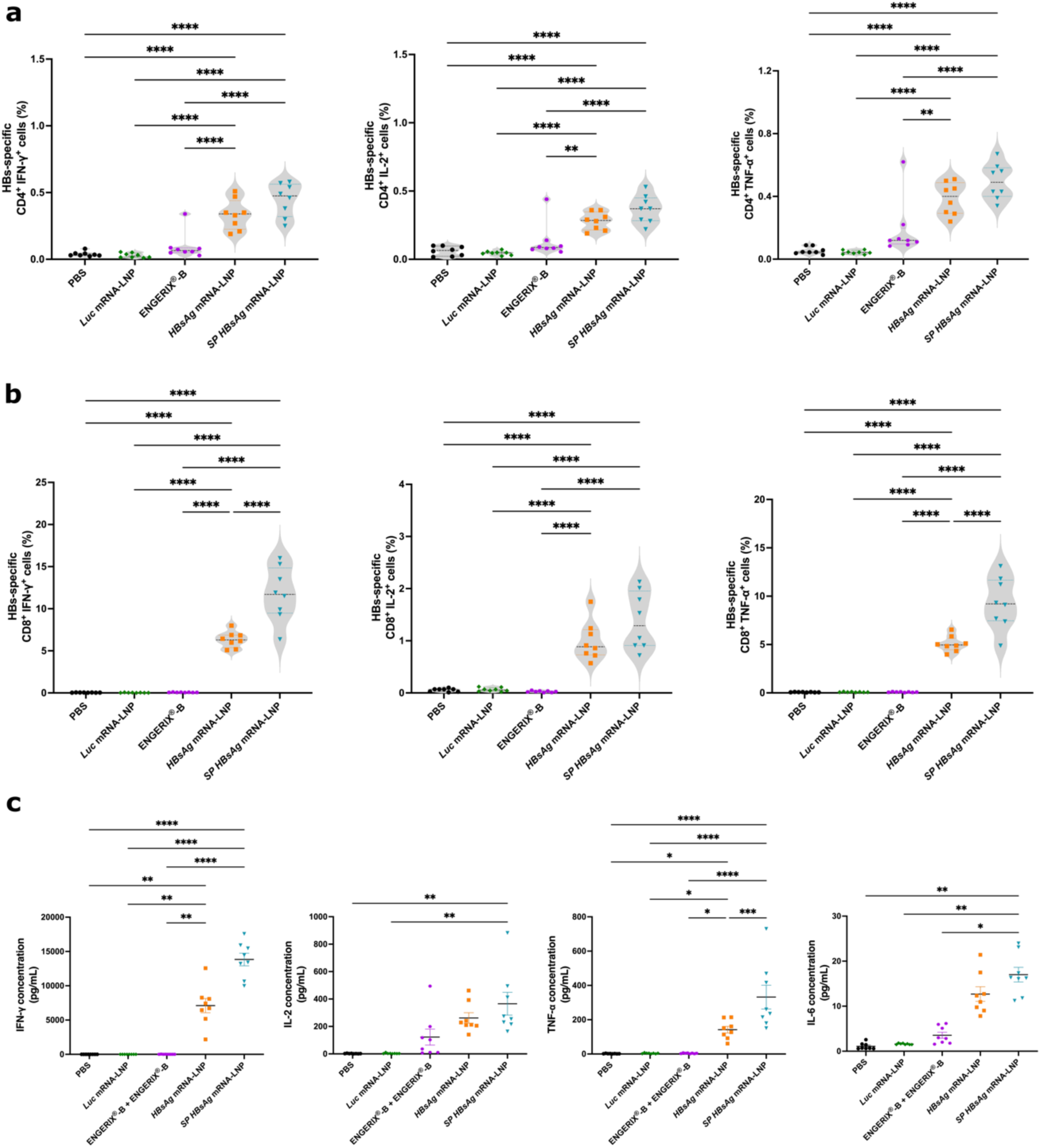
*HBsAg* mRNA-LNP vaccines induce robust antigen-specific CD4^+^ and CD8^+^ immune responses. (a) Percentages of IFN-γ^+^, IL-2^+^ and TNF-α^+^-producing antigen-specific CD4^+^ and (b) CD8^+^ T cells from their respective previous CD4^+^ and CD8^+^ T cell population after peptide stimulation of isolated splenocytes. (c) IFN-γ, IL-2, TNF-α, and IL-6 secreted by spleen cells were quantified in cell culture supernate 24 hours after HBsAg-peptide stimulation. Data are given as the means ± SEM for animal cohorts (8 mice/group). Significant differences tested compared to the other groups: *p <0.05; **p <0.01; ***p <0.001; ****p <0.0001 (one-way ANOVA, Tukey’s multiple comparisons test).

Cytokines in the supernates obtained from splenocytes cultured and stimulated with peptides derived from HBsAg for 24 hours were quantified. Relative to splenocytes derived from control mice that had received either PBS, *Luc* mRNA-LNP, or ENGERIX^®^-B, splenocytes derived from all mouse cohorts immunized with mRNA-LNP vaccines produced high levels of IFN-γ, IL-2, TNF-α, and IL-6 (Figure 3c). Notably, the incorporation of SP into the N-terminus of *HBsAg* mRNA resulted in enhanced production of cytokines when compared with the levels secreted by splenocytes derived from *HBsAg* mRNA-LNP vaccinated mice.

### *HBsAg* mRNA-LNP vaccination boosts ENGERIX^®^-B vaccine-induced immune responses

While many countries adhere to the introduction of anti-HBV immunization during the first 24 hours of life, frequently, the completion rate for the HBV vaccination schedule is low due to different reasons, such as low socio-economic status, poor education, and difficulties in accessing healthcare facilities, among others. ^31–33^ Additionally, 5% of vaccinated individuals do not mount immunity against the virus.^34^ In this regard, it becomes relevant to evaluate whether the mRNA-based vaccines studied here can boost ENGERIX^®^-B-induced immune response. Immunization schemes were executed as in previous studies (prime-boost vaccination 28 days apart). *HBsAg* mRNA-LNP and *SP-HBsAg* mRNA-LNP vaccines were used only to boost protein-based priming (ENGERIX^®^-B), while complete ENGERIX^®^-B immunization (prime-boost) was used as a control (Figure 4a). Notably, boosting with either *HBsAg* mRNA-LNP or *SP-HBsAg* mRNA-LNP significantly increased antigen-specific endpoint IgG levels compared to the control group vaccinated with ENGERIX^®^-B only (Figure 4b). IgG2c/IgG1 ratios were not significantly different between the groups (Figure S4). Furthermore, the mRNA-LNP-boosted groups improved the cellular immune response. This effect was particularly evident when ENGERIX^®^-B priming was boosted with *SP-HBsAg* mRNA-LNP; a substantial increase in cytokine-producing CD4^+^ and CD8^+^ T cells was detected (Figure 4c and 4d). Spleen cells derived from mRNA-LNP-boosted mice, particularly *SP-HBsAg* mRNA-LNP-boosted, secreted high levels of IFN-γ, IL-2, TNF-α, and IL-6 after HBsAg-peptides stimulation (Figure 4e).

**Figure 4.**
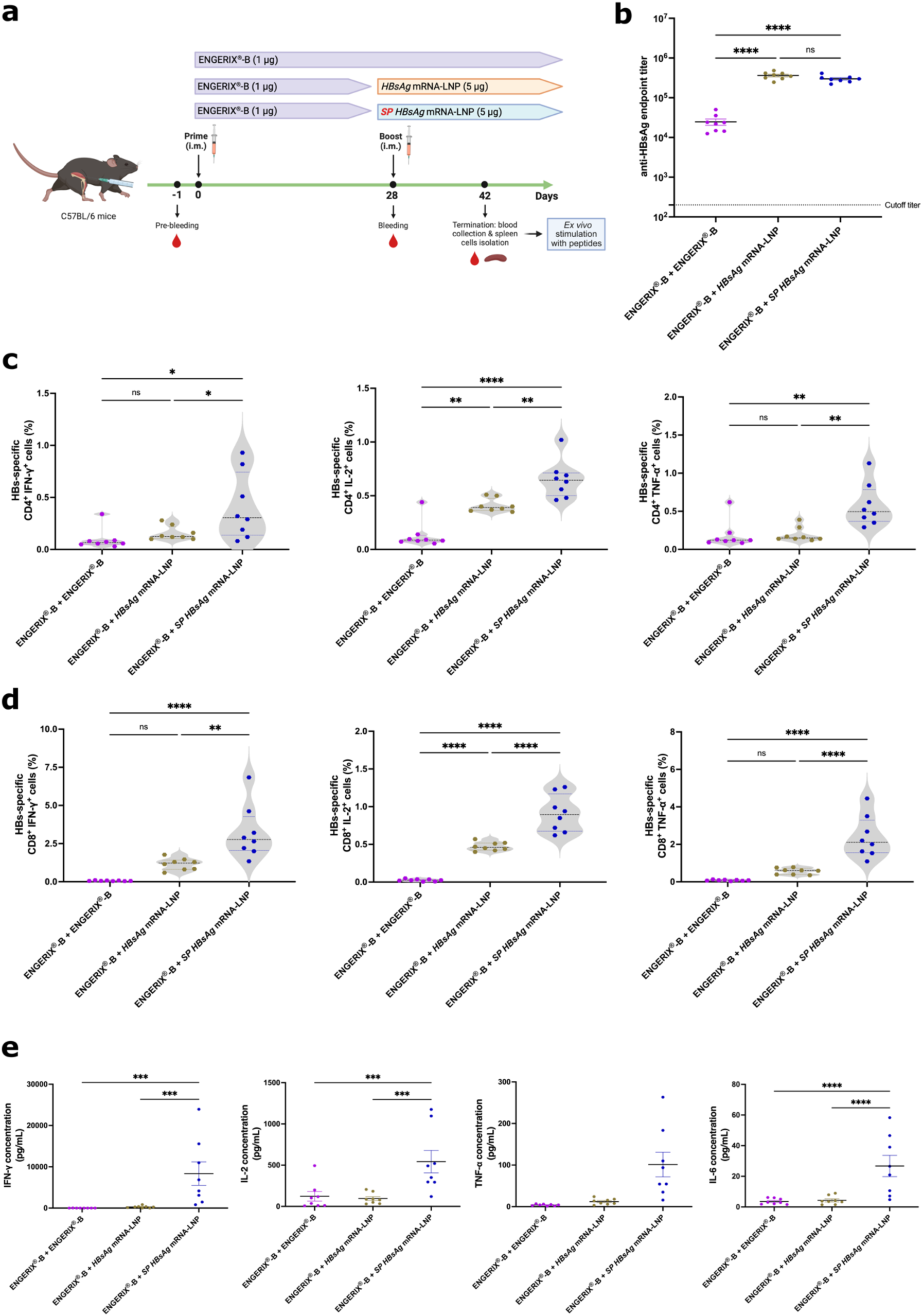
Heterologous (mRNA-protein) immunization enhances immune responses compared with homologous protein immunization with the commercial ENGERIX^®^-B vaccine. (a) Representation of the heterologous immunization scheme: 3 experimental cohorts (each n=8) were primed i.m. at day 0 with ENGERIX^®^-B. On day 28 groups were immunized as follows: (i) 1 μg ENGERIX^®^-B vaccine, (ii) 5 μg *HBsAg* mRNA-LNP, and (iii) 5 μg *SP-HBsAg* mRNA-LNP. (b) Total anti-HBsAg IgG titers at termination of the immunization schedule (d28). (c) Percentages of IFN-γ^+^, IL-2^+^ and TNF-α^+^-producing antigen-specific CD4^+^ and (d) CD8^+^ T cells from their respective previous CD4^+^ and CD8^+^ T cell population after peptide stimulation of cell isolates from spleen. (e) Secreted cytokine concentration (IFN-γ, IL-2, TNF-α, and IL-6) from spleen cells analyzed 24 hours after peptide stimulation. Data are the means ± SEM for animal cohorts (8 mice/group). Significance tested compared to the other groups: no significant differences (ns); *p < 0.05; **p < 0.01; ***p < 0.001; ****p < 0.0001 (one-way ANOVA, Tukey’s multiple comparisons test).

### HBsAg mRNA-LNP vaccines induce protective immunity in mice challenged with HBV

Immunocompetent mice were immunized two times 28 days apart (5 μg dose) with either *HBsAg* mRNA-LNP or *SP-HBsAg* mRNA-LNP. Twenty-eight days after administration of the boost, animals were challenged by infection with a recombinant adeno-associated virus subtype-8 modified with 1.3 copies of the HBV genome (rAAV8-1.3HBV) (Figure 5a). AAV8 is highly hepatotropic, therefore, the rAAV8-1.3HBV system has been widely used to study the anti-viral effects of different drug product candidates against HBV.^35^ More importantly, rAAV8-1.3HBV allows an efficient and persistent HBV infection characterized by fast and long-lasting serum HBsAg levels.^36^ Therefore, this model can mimic a chronic HBV infection, and could also be used to represent a strong acute HBV infection. All animals preserved clinical health during the study as their body weight was not negatively affected (Figure S5a). In the current study, plasma anti-HBsAg antibodies quantified at the experimental endpoint revealed high IgG titers in vaccinated but not in PBS-injected control animals (Figure 5b). One and two weeks after rAAV8-1.3HBV infection, high levels of HBsAg protein were detected in the plasma of the control cohort but were undetectable in mice immunized with the mRNA-LNP vaccines (Figure 5c).

**Figure 5.**
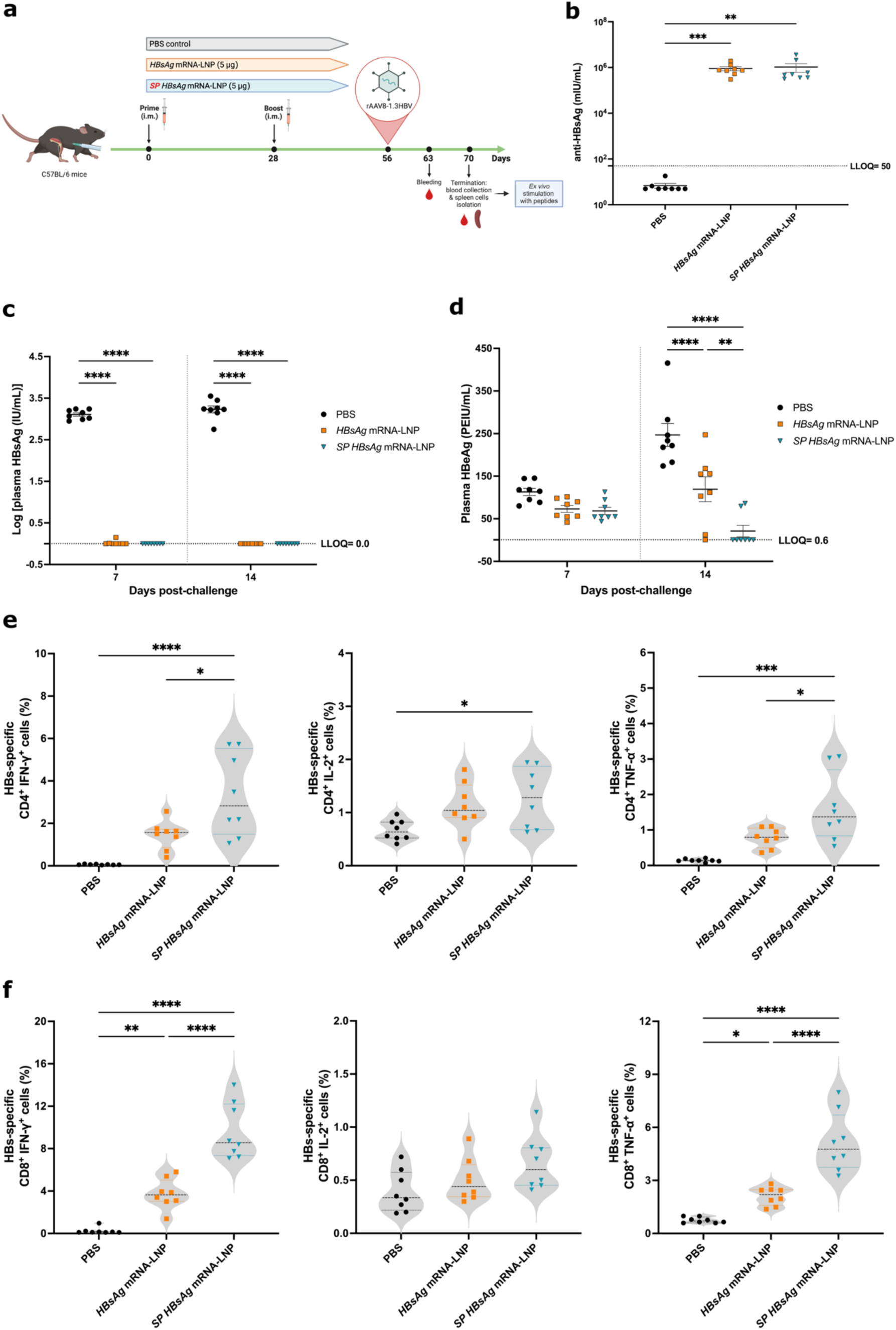
Optimized mRNA-LNP vaccines prevent HBV replication in rAAV8-1.3HBV challenged mice. (a) Experimental schedule. Mice were immunized with mRNA-LNP vaccines on day 0 (prime). Four weeks after priming, the second dose (boost) was injected (d28). PBS was used as negative control. On day 56, animals were challenged with AAV/HBV viral genome. The experiment was finalized on day 70, and blood and spleen were collected for further analyses. Blood was also collected on day 63. (b) Total anti-HBsAg IgG concentration at termination of the immunization schedule (d70). (c) Measurement of plasma HBsAg and (d) HBeAg on day 7 and day 14 post-challenge. (e) Percentages of peptide-stimulated IFN-γ^+^, IL-2^+^ and TNF-α^+^-producing antigen-specific CD4^+^ and (f) CD8^+^ T cells quantified after overnight culture. Data are the means ± SEM for animal cohorts (8 mice/group). Significance tested compared to the other groups: *p < 0.05; **p <0.01; ***p <0.001; ****p <0.0001 (one-way ANOVA, Dunn’s and Tukey’s multiple comparisons test; two-way ANOVA, Tukey’s multiple comparisons test). LLOQ: lower limit of quantification.

Plasma Hepatitis B “e” antigen (HBeAg) levels, another indicator of HBV infection, were also measured at one and two weeks post-infection (Figure 5d). HBeAg was detected in all experimental cohorts 1 week after infection; significant differences were not observed between groups. Two weeks post-challenge, HBeAg levels were further increased in the control group but remained unaltered in *HBsAg* mRNA-LNP-vaccinated mice. Interestingly, HBeAg levels in *SP-HBsAg* mRNA-LNP-vaccinated mice (6 out of 8 animals) were below the lower limit of quantification (LLOQ). The specific cellular immune responses were examined at the experimental endpoint. Immunization with mRNA-LNPs increased the splenic IFN-γ^+^, IL-2^+^, and TNF-α^+^ CD4^+^ (Figure 5e) and CD8^+^ (Figure 5f) T cell populations significantly: the largest increases were observed in the group vaccinated with *SP-HBsAg* mRNA-LNP.

### *HBsAg* mRNA-LNP vaccination suppresses the viral load and overcome immune tolerance in HBV-carrier mice

Among the mouse models used to represent CHB, the AAV-mediated transfection mouse model is based on viral transduction of the HBV genome, initiating HBV replication and secretion of infectious HBV virions.^35^ The viral construct used in this work (rAAV8-1.3HBV) was injected intravenously (i.v.) into C57BL/6 mice to establish the CHB mouse model (HBV-carrier mice). HBV-carrier mice are characterized by persistent viral replication with long-lasting HBsAg levels in serum. Moreover, these mice were reported to have T-cell immune tolerance and to be completely resistant to immunization with a conventional aluminum-adjuvanted vaccine.^36^ To overcome this CHB-associated immune tolerance, HBV-carrier mice were primed and then boosted twice with mRNA-LNP vaccines. As depicted in Figure 6a, experimental cohorts were organized in four groups: (i) control, injected with PBS, (ii) 10 μg *HBsAg* mRNA-LNP, (iii) 10 μg *SP-HBsAg* mRNA-LNP, and (iv) a combinatorial approach comprising priming with 10 μg *HBsAg* mRNA-LNP, followed by first boost with a mix of 5 μg *HBsAg* mRNA-LNP and 5 μg *SP-HBsAg* mRNA-LNP, and finally a second boost with 10 μg *SP-HBsAg* mRNA-LNP. Considering that the long-term chronic HBV infection is characterized by a tolerogenic microenvironment with a weak virus-specific immune response and an overall impairment in the CD8^+^ T cell responses, the combinatorial approach with both vaccine candidates aimed to prime the CD4^+^ T cell responses to improve collaborative communication with the non-responsive CD8^+^ T cells. Therefore, a prime immunization with the *HBsAg* mRNA-LNP vaccine was performed as this construct induced a more equilibrated humoral response in favor of IgG2c/IgG1 ratio. Next, a first boost immunization combining both *HBsAg* as well as *SP-HBsAg* mRNA-LNP was administered, followed by a second boost with the *SP-HBsAg* mRNA-LNP candidate, as a potent CD8^+^ T cell response activator.

**Figure 6.**
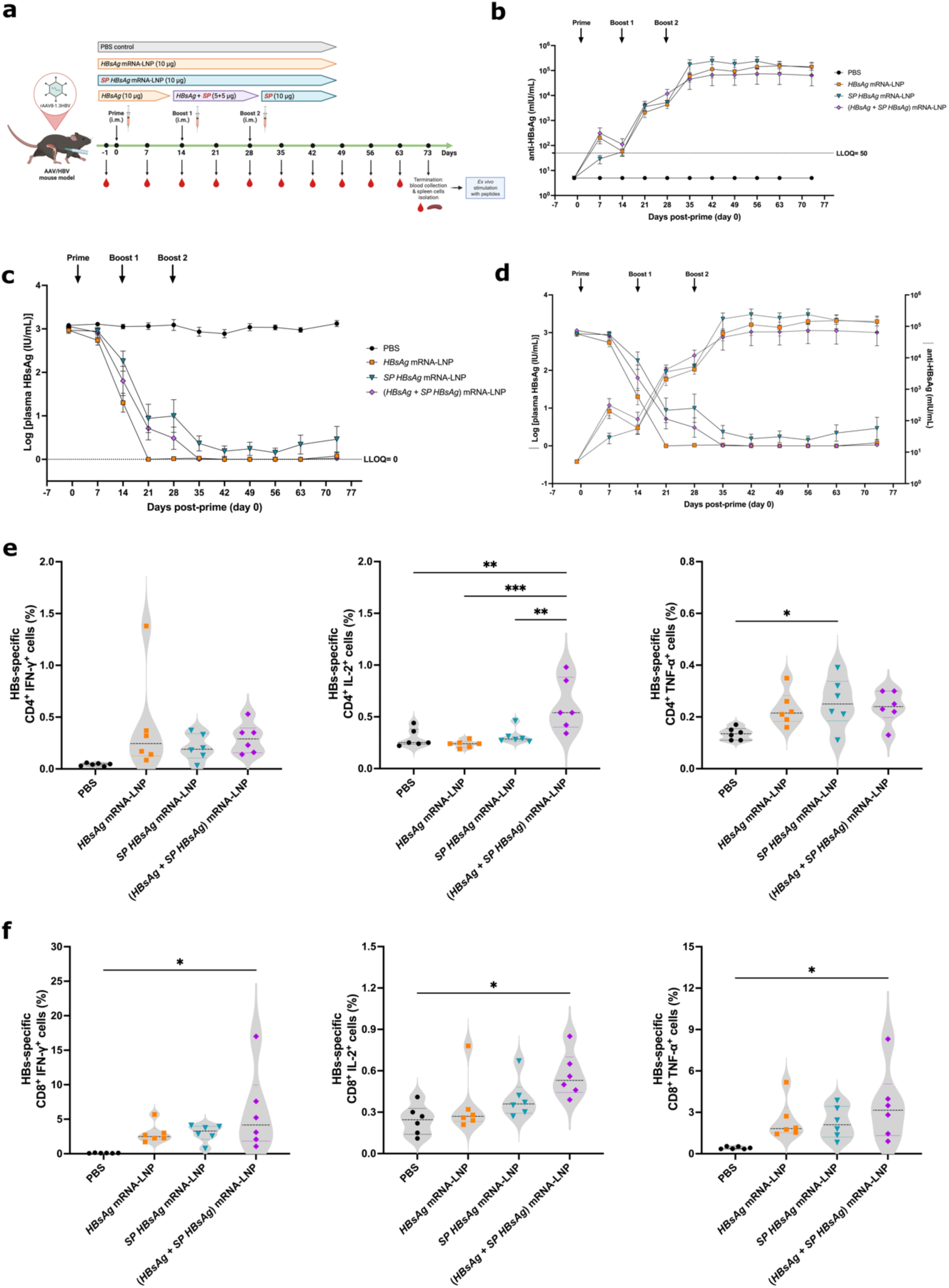
*HBsAg* mRNA-LNP vaccination reduces serum HBsAg and induces cellular immunity in CHB mice. (a) Experimental design. Mice were injected i.v. with AAV/HBV viral genome on 28 days before prime (CHB mice). CHB mice were vaccinated i.m. with (i) 10 μg *HBsAg* mRNA-LNP, (ii) 10 μg *SP-HBsAg* mRNA-LNP on days 0 (Prime), 14 (Boost 1) and 28 (Boost 2), and (iii) 10 μg *HBsAg* mRNA-LNP on day 0 (Prime), 5 μg *HBsAg* mRNA-LNP + 5 μg *SP-HBsAg* mRNA-LNP on day 14 (Boost 1) and 10 μg *SP-HBsAg* mRNA-LNP on day 28 (Boost 2). PBS was used as negative control. The experiment was terminated on day 73, and blood and spleens were collected for further analyses. Blood was also collected on day -1, and days 7, 14, 21, 28, 35, 42, 49, 56 and 63 post-prime. (b) Total anti-HBsAg IgG concentration determined at all experimental points. (c) Measurement of plasma HBsAg and (d) reverse correlation of serum anti-HBsAg antibody and plasma HBsAg concentrations over time. (e) Percentages of IFN-γ^+^, IL-2^+^ and TNF-α^+^-producing antigen-specific CD4^+^ and (f) CD8^+^ T cells quantified after overnight stimulation with peptides. Data are the means ± SEM for animal cohorts (6 mice/group). Significantly different from other groups: *p <0.05; **p <0.01; ***p <0.001 (one-way ANOVA, Tukey’s multiple comparisons test). LLOQ: lower limit of quantitation.

The body weight of the HBV-carrier animals after immunization developed normally (Figure S5b), and seroconversion was observed in all vaccinated animals. The HBV-carrier mice that were primed with the *HBsAg* mRNA-LNP vaccine (groups ii and iv) exhibited a faster increase in anti-HBsAg antibody levels. A plateau in antibody concentrations was reached after the second boost (Figure 6b). Correspondingly, plasma HBsAg were almost undetectable after the second boost in all vaccinated groups (Figure 6c and 6d). Noteworthy, HBsAg levels (indicating viral infection) in mice immunized only with *SP-HBsAg* mRNA-LNP vaccine remained low but above the LLOQ during the 73-day study (Figure 6c). Interestingly, the group vaccinated with a combination of *HBsAg* mRNA-LNP and *SP-HBsAg* mRNA-LNP exhibited an improvement as reflected by lowered HBsAg plasma levels when comparing it with the individual *SP-HBsAg* group. mRNA-LNP vaccines elicited strong CD4^+^ and CD8^+^ T cell responses by expressing anti-viral cytokines such as IFN-γ, IL-2, and TNF-α in response to HBsAg peptide stimulation (Figures 6e-f). Notably, the combinatorial immunization strategy generated the highest frequency of cytokine-secreting CD8^+^ T cells, outperforming the cellular responses observed in those groups that received the individual candidate vaccines at the same overall dose of 10 μg. In summary, the data demonstrate that both vaccine candidates were effective in a mouse model of CHB. Seroconversion was successful, independent of the incorporation of the signal peptide into the antigenic sequence. An improvement in the cell-mediated immunity by the presence of the signal peptide was only observed in the combinatorial approach, indicating a greater restoration of immune control.

## Discussion

mRNA-based vaccine platforms possess advantages over conventional ones (live-attenuated pathogen, inactivated pathogen, and pathogen-specific protein subunit) due to their ease of production, flexibility and scalability.^20^ Additionally, mRNA-LNP vaccines do not require the addition of adjuvants since both the mRNA molecule and LNP component are inherently able to activate innate immune cells.^37^ Th1-polarized immune responses, which generally promote anti-viral immunity, often occur in response to mRNA-based vaccination.^37,38^

In this study, HBsAg was selected as the vaccine target since it is highly immunogenic. Moreover, HBsAg-specific seroconversion and decreases in HBsAg serum levels achieved in some CHB patients are often associated with functional cure. In the present study, a nucleoside-modified mRNA construct that encodes the HBsAg protein sequence (i.e., *HBsAg* mRNA) was synthesized, and hmoDCs were transfected as an immunologically relevant cell type to characterize *HBsAg* mRNA expression. Transfection with *HBsAg* mRNA induced HBsAg protein production. Flow cytometric analysis demonstrated the presence of HBsAg both intracellularly and on the surface of *HBsAg* mRNA-transfected hmoDCs. Collectively, these findings demonstrate the ability of the *HBsAg* mRNA to transfect immunologically relevant cells and to induce the production of HBsAg protein.

Optimal T cell priming occurs when: 1) antigenic peptide sequences are presented in association with MHC molecules, 2) costimulatory molecule expression on the surface of APCs is up-regulated, and 3) secretion of proinflammatory cytokines occurs. In the experiments reported here, *HBsAg* mRNA significantly increased the cell-surface expression of CD80, CD86, HLA-DR, and CD40 by transfected hmoDCs. *HBsAg* mRNA transfection also induced IP-10 (CXCL10; an IFN-γ responsive gene), and TNF-α secretion by hmoDCs.^38^

The exact contribution of the nucleoside-modified and purified mRNA and the LNP components to the adjuvanticity of mRNA-LNP vaccines is not fully understood.^37^ In the current study, the enhanced expression of cell-surface costimulatory molecules and cytokine secretion occurred only after hmoDCs were transfected with *HBsAg* mRNA, but not with *ova* mRNA. This observation indicates, in principle, that under our *in vitro* experimental conditions the nucleoside-modified mRNA alone does not account for any specific adjuvant effect on hmoDCs. It is relevant to note, however, that HBsAg protein alone also exerts a self-adjuvant effect on hmoDCs *in vitro*, upregulating the expression of CD80, CD83, CD86, and HLA-DR.^39^ Further investigations need to be carried out to better understand the biological causes of these self-adjuvancy of HBsAg. Conceivably, it may be the combined effects of *HBsAg* mRNA and the translated protein product that stimulate the innate immune response of hmoDCs observed in this work.

The early immune effects of *HBsAg* mRNA-LNP injected i.m. into mice were assessed 24 hours after administration. Both the DCs and T cells in the spleen and inguinal lymph nodes were effectively activated, indicating that immunization with *HBsAg* mRNA-LNP generated a protein product (HBsAg) in a setting that was optimal for the induction of adaptive B and T cell responses.

To improve antigen presentation, an MHC class I signal peptide (SP) was incorporated into the *HBsAg* mRNA construct. Next, mice were immunized i.m. with two doses of mRNA vaccines or the licensed anti-hepatitis B vaccine, ENGERIX^®^-B. Quantitation of anti-HBsAg titers in immunized mice demonstrated a robust humoral response to *HBsAg* and *SP-HBsAg* mRNA-LNP vaccines. Incorporating the N-terminal signal peptide did not negatively impact the production of antibodies induced by the *HBsAg* mRNA vaccine; both mRNA vaccine constructs significantly outperformed the response elicited by the protein-based adjuvanted vaccine, ENGERIX^®^-B. In this regard, other investigators report that mRNA-LNP vaccines promoted increases in T follicular helper cells, formation of germinal centers, B cell activation, and the production of high-avidity antibodies.^38,40^ Consequently, studies that compare vaccine preparations often report that mRNA-LNP vaccines outperform adjuvanted protein vaccines.^41,42^

Seroconversion is not sufficient to overcome CHB infection^43–45^. A Th1-, rather than a Th2-type immune response is preferred for generating effector T cell-mediated immunity. IgG2c/IgG1 ratio is a reliable parameter for determining polarization towards Th1 or Th2. In the current study, the *SP-HBsAg* mRNA vaccine construct induced the production of large amounts of IgG2c antibodies. High IgG2c/IgG1 ratios indicate polarization towards Th1-type responses. IgG2c antibodies possess a high capacity to bind to all activating Fcγ-receptor (FcγRs), thus strongly promoting immunity.^46,47^ Interestingly, increases in IgG2c are often observed in adjuvanted vaccines.^48–50^ Those mice immunized with *HBsAg* mRNA-LNP showed a more balanced IgG response. In contrast, mice immunized with the protein-based vaccine, ENGERIX^®^-B, failed to exhibit increases in the IgG2c/IgG1 ratio. This was evidenced by the predominance of IgG1-type antibodies in serum, whereas IgG2c titers were negligible. ENGERIX^®^-B is adjuvanted with alum, which is widely known to elicit Th2 immune responses and the production of antigen-specific antibodies; it is incapable, however, of stimulating Th1 or cytotoxic T cell responses.^51^

HBV-specific CD4^+^ and CD8^+^ T cell responses are essential for virus clearance and control.^43^ Additionally, secreted antiviral cytokines can interfere with cccDNA activity and induce its degradation.^44^ Indeed, mRNA-LNP vaccines often elicit CD4^+^ and CD8^+^ T cells that express Th1 cytokines.^42,52–56^ Humans immunized with COVID-19 mRNA vaccines, for example, exhibit long-lasting antigen-specific CD4^+^ T cells and IFN-γ-producing CD8^+^ T cells.^37,57,58^ Vaccination of mice with *HBsAg* and *SP-HBsAg* mRNA-LNP in the current study induced HBsAg-specific IFN-γ, IL-2 and TNF-α producing CD4^+^ and CD8^+^ T cells. The elevated production of IFN-γ, TNF-α and IL-2 in response to HBsAg-derived peptide stimulation is consistent with the Th1-type responses often seen following the administration of mRNA vaccines.^41^ Interestingly, the N-terminal modification using the ER translocation signal from MHC class I resulted in the highest cytokine production by CD8^+^ and CD4^+^ T cells. Considering this, as well as its ability to promote the highest IgG2c/IgG1 ratio, the *SP-HBsAg* mRNA-LNP vaccine constitutes the candidate capable of mounting the strongest specific immune cellular response to HBV.

mRNA-based vaccination could be potentially introduced as a booster option in those patients whose vaccination scheme has been initiated but not completed or in the fraction of vaccine non-responders.^34,59,60^ To address this question, mice were primed with ENGERIX^®^-B and boosted with mRNA-based vaccines. Animals generated antigen-specific antibody titers that were comparable to those achieved following an mRNA-LNP prime-boost scheme. Interestingly, despite failing to induce IgG2c antibodies, heterologous (protein prime-mRNA boost) vaccination induced CD4^+^ and CD8^+^ T cell responses that were comparable to those yielded by homologous mRNA-LNP vaccination. These results are consistent with a previous study, in which priming with a protein-based vaccine followed by an mRNA vaccine boost led to a robust CD8^+^ T cell response.^61^

HBsAg detected in sera is generally recognized as a marker for persistent HBV infections.^62^ To evaluate the protective efficacy of the vaccines, mice vaccinated with either *HBsAg* mRNA-LNP or *SP-HBsAg* mRNA-LNP were challenged by infection with rAAV8-1.3HBV. Both vaccine candidates mounted humoral and cell-mediated immune responses that prevented increases in plasma HBsAg levels.

Perinatal transmission of HBV is often associated with unvaccinated pregnant women who are positive for the HBeAg; transmission is significantly less frequent when the mothers are HBeAg-negative.^63^ Besides being a marker for HBV infection, HBeAg is the only antigenic protein capable of trespassing the placental barrier. Consequently, HBeAg has been described as an immunomodulator and a tolerogenic protein.^64^ Although the role of HBeAg in chronicity is unclear, the repercussions of its tolerogenic effect on immature immune systems undoubtedly contribute to the development of CHB in newborns.^63,65^ Relevant to the current study, high HBeAg levels were found within 7 days after the challenge in the three vaccinated groups. Notably, vaccination with *SP-HBsAg* mRNA-LNP and, to a lesser extent, *HBsAg-*mRNA-LNP successfully reduced HBeAg to baseline levels 14 days after HBV challenge.

To assess if our anti-HBV mRNA-LNP vaccine candidates can protect from chronic infection, HBV-carrier mice mimicking CHB infection were immunized. Regardless of HBV-carrier mice were vaccinated with *HBsAg* mRNA-LNP, *SP-HBsAg* mRNA-LNP, or a combination of them, a strong immune response was induced after the three immunizations. Remarkably, seroconversion and clearance of HBsAg in plasma were achieved and maintained at least for 73 days post-prime immunization. Reversing immune tolerance alone fails to restore antibody production, therefore, virus-specific CD8^+^ and CD4^+^ T cell activation must accompany seroconversion for successful viral clearance.^43,66,67^ Of particular interest, we observed robust cell-mediated immune response in the combination group, where an improvement in HBsAg-specific CD8^+^ T cell responses was observed compared to the individual vaccination schemes.

In summary, the work described here establishes optimized nucleoside-modified mRNA-LNP vaccines and immunization schemes against hepatitis B for use both therapeutically, as well as prophylactically. In contrast to the licensed ENGERIX^®^-B vaccine, immunization with mRNA-LNP vaccines induced robust T cell responses, as well as elevated HBsAg-specific antibody titers. Additionally, the study highlights the improved, Th-1 biased, immune responses resulting from incorporating an SP signal into the mRNA sequence. The ability of the mRNA vaccines to induce both strong humoral- and cell-mediated immune responses provides effective protection against viral challenges in a mouse model. We also demonstrated that ENGERIX-induced immune responses can be robustly boosted with HBsAg mRNA-LNPs. Further, the tested vaccination schemes with HBsAg mRNA-LNP vaccines were therapeutically effective in overcoming HBV infection in a CHB mouse model. As such, HBsAg mRNA-LNP vaccines have the potential to eradicate HBV in chronically infected patients. Overall, these data encourage the continued evaluation of HBsAg mRNA-LNP vaccines as potential therapeutic candidates to cure CHB.

## Methods

### Cell line

HepG2 cells were purchased from The American Type Culture Collection (ATCC; Manassas, VA, USA), ATCC^®^ Number: HB-8065^TM^. HepG2 cells were cultured in RPMI-1640 medium supplemented with 10% heat-inactivated fetal bovine serum (iFBS), 100 U/mL of penicillin, 100 μg/mL streptomycin, 1% L-glutamine and 5 mM HEPES.

### Animals

Eight- to 10-week-old wild-type C57BL/6Ncr mice were purchased from Charles River Laboratories. Five mice/cage were housed in a dedicated vivarium (18-23°C ambient temperature, 40-60% humidity, 14 h:10 h light cycle) with clean food, water and bedding. Experimental cohorts were composed of both males and females selected at random; randomized littermate controls were included in each experiment. All animal procedures were approved by the local authorities (Landesuntersuchungsamt Rhineland-Palatinate). Ethical approval was granted by the Landesuntersuchungsamt LUA, Koblenz, AK G 23-1-006.

Five-week-old C57BL/6 mice obtained from the Shanghai Laboratory Animal Center of Chinese Academy of Sciences were used for challenge and chronic hepatitis B experiments. HBV-carrier C57BL/6 mice (AAV/HBV mouse model) were generated by infecting them with rAAV8-1.3HBV (Ayw, D type) four weeks prior to vaccination. The studies were approved by the WuXi Institutional Animal Care and Use Committee (protocol ID01-013-2021v1.0) and conducted by WuXi AppTec (Shanghai) Co., Ltd.

### mRNA-LNP vaccine production

mRNA constructs were generated using the sequence that encodes hepatitis B, genotype A surface antigen (*HBsAg* mRNA) (GenBank: BAD91275.1). For *in vitro* tests and *in vivo* early activation, an *HBsAg* mRNA construct from TriLink Biotechnologies (San Diego, USA). Briefly, the construct was generated by using a proprietary vector that contains a T7 promoter, optimized 5’ UTR with a strong kozak sequence, a 3’ UTR derived from a mouse alpha-globin gene, and a 120 nucleotide-long poly(A) tail. For early activation *in vivo* experiments (Figure S1), the *HBsAg* mRNA was encapsulated in LNP using a NanoAssemblr Ignite™ device for manufacturing the formulation (Precision Nanosystems, Vancouver, Canada). Genvoy-ILM™ lipid mix (Precision Nanosystems, Vancouver, Canada), containing 50% ionizable lipid, 10% DSPC, 37.5% cholesterol, and 2.5% PEG-lipid was mixed with mRNA in 50 mM Acetate buffer, pH 4.0, using an N/P ratio of 6. The mean hydrodynamic size and distribution (PDI) of the LNP formulation was measured by dynamic light scattering (DLS) in a Nano ZS Zetasizer (Malvern Instruments Corp., Malvern, UK) with a mean diameter in the range of 80-90 nm with low PDI (<0.05). HBsAg mRNA encapsulation efficiency of ∼95% was achieved as determined using a Ribogreen assay (Thermo Fisher Scientific, Waltham, MA).

To enhance HBsAg-specific immune responses, mRNA constructs were optimized by incorporating an MHC class I signal peptide (SP) in the 5’ region and a transmembrane and cytosolic trafficking domain of MHC class I (MITD) in the 3’ region flanking the *HBsAg* mRNA sequence. As a result, two new constructs were obtained: *SP-HBsAg* mRNA and *SP-HBsAg-MITD* mRNA. The mRNA vaccines using these sequences were made as previously described.^68,69^ Briefly, the codon-optimized genes were synthesized (Genscript). Constructs were ligated into mRNA production vectors, vectors were linearized, and a T7-driven in vitro transcription reaction (Megascript, Ambion) was performed to generate mRNA with 101 nucleotide-long poly(A) tails. Capping of mRNA was performed in concert with transcription through addition of a trinucleotide cap1 analog, CleanCap (TriLink), and m1Ψ-5’-triphosphate (TriLink) was incorporated into the reaction instead of UTP. Cellulose-based purification of mRNA was performed as described.^68^ mRNAs were then assessed on an agarose gel before storing at −20°C.

Cellulose-purified *HBsAg, SP-HBsAg* and *SP-HBsAg-MITD* mRNAs were encapsulated in LNPs using a self-assembly process in which an aqueous solution of mRNAs at pH 4.0 is rapidly mixed with a solution of lipids dissolved in ethanol. LNPs used in this study, consisted of a mixture of the ionizable cationic lipid (pKa in the range of 6.0–6.5, proprietary to Acuitas Therapeutics)/phosphatidylcholine/cholesterol/PEG-lipid (50:10:38.5:1.5 mol/mol) as described in the patent application WO 2017/004143. The average hydrodynamic diameter was ∼80 nm with a polydispersity index of 0.02–0.06 as measured by dynamic light scattering using a Zetasizer Nano ZS (Malvern Instruments Ltd, Worcestershire, UK) and an encapsulation efficiency of ∼95% was achieved as determined using a Ribogreen assay (Thermo Fisher Scientific, Waltham, MA).

### Human monocyte-derived dendritic cells (hmoDCs)

Blood samples were obtained from healthy volunteers who donated at the blood bank (University Medical Center Mainz, Germany). All donations were collected upon informed consent. Peripheral blood mononuclear cells (PBMCs) were enriched and collected by density gradient centrifugation using Histopaque-1077 (Sigma-Aldrich, Steinheim am Albuch, Germany). The monocytes were isolated from PBMCs by positive selection using a CD14 Microbeads kit (Miltenyi Biotec, Bergisch Gladbach, Germany) according to the manufacturer’s protocol, and seeded into six-well suspension culture plates at a density of 10^6^ cells/mL of serum-free X-VIVO 15 medium (Lonza, Walkersville, MD) supplemented with 1% penicillin-streptomycin, 25 ng/mL of recombinant human IL-4 (ImmunoTools GmbH, Friesoythe, Germany) and 100 ng/mL of recombinant human granulocyte macrophage colony stimulating factor (ImmunoTools). The cells were incubated for 4 days at 37°C in a humidified environment containing 5% CO_2_; half the spent culture medium was replaced with fresh supplemented X-VIVO 15 medium on day 2.5.

### Transfection of immature hmoDCs and HepG2 cells

The suspension cells that in large part constitute immature hmoDCs after 4 days of culture were collected and seeded into 24-well tissue culture plates at 1.2 x 10^6^ cells/mL of supplemented X-VIVO 15 medium. After an additional 24 h incubation, the cells were transfected with 500 ng *HBsAg* mRNA mixed with 1 μL Lipofectamine MessengerMax (ThermoFisher) in accordance with the manufacturer’s instructions. The hmoDCs transfected with 500 ng *ova* mRNA mixed with 1 μL Lipofectamine MessengerMax served as a negative mRNA control. Recombinant HBsAg (Hytest, Turku, Finland) or ENGERIX^®^-B, anti-hepatitis B vaccine (GlaxoSmithKline Biologicals s.a., GSK; Rixensart, Belgium) added to the culture medium at 1 μg/mL served as the HBsAg protein positive controls. The addition of Lipofectamine MessengerMax alone served as a negative control. The results were compared to untreated hmoDCs derived from the same donor. Cells and cell culture supernatants were analysed 24 h after transfection or stimulation.

HepG2 cells were seeded into 24-well tissue culture plates at 5 x 10^5^ cells/mL in RPMI-1640 culture medium. After 24 h incubation, the cells were transfected with 500 ng of mRNA constructs mixed with 1 μL Lipofectamine MessengerMax (ThermoFisher) according to the manufacturer’s instructions. Non-transfected HepG2 cells were used as negative control.

### Intracellular (ICS) and extracellular (ECS) staining

The hmoDCs transfected with *HBsAg* mRNA or incubated with HBsAg protein (positive control) were subjected to ICS and ECS to characterize the expression of *HBsAg* mRNA by the antigen-presenting cells. Twenty-four hours after transfection, the cells were detached from the substratum by incubating with 1X DPBS (2 mM EDTA and 0.5% BSA) for 30 min on ice, then washed with fluorescence-activated cell sorting (FACS) buffer (1X DPBS containing 2% iFBS). Subsequently, the cells were incubated at 4°C for 15 min with 10% Privigen® Immunoglobulin solution (CSL Behring; Marburg, Germany), an Fc receptor blocker.

For ECS, hmoDCs were incubated with biotin-conjugated anti-HBsAg antibody (ab68520; Abcam, Cambridge, United Kingdom) at 4°C for 30 min, washed, then incubated with the LEGENDplex SA-PE (BioLegend; San Diego, CA, USA) for 30 min at room temperature in the dark and washed twice. For ICS, the cells were fixed and permeabilized using Cytofix/Cytoperm fixation/permeabilization kit (BD Biosciences; Franklin, NJ, USA) according to the manufacturer’s instructions. The cells were then incubated with biotin-conjugated anti-HBsAg antibody (Abcam), washed, incubated with LEGENDplex SA-PE (BioLegend), and washed twice. Stained cells were acquired with an LSR II flow cytometer (BD Biosciences) and analyzed with FlowJo software version 10.8 (Ashland, OR; Becton, Dickinson and Company).

### Western blot analysis

HepG2 cells were transfected with the mRNA constructs. The cells were detached from the substratum after 24 hours by incubating with trypsin for 10 min at 37°C. The cells were collected and centrifuged; the cell pellet was resuspended in 40 μL lysis buffer (Pierce^TM^ IP lysis buffer, 1X protease inhibitor cocktail and 1 mM DTT) and incubated for 30 min on ice. Total lysate protein was quantified using Qubit^TM^ Protein Assay Kit (ThermoFisher). SDS-PAGE was performed using equal protein concentrations per sample. Subsequently, the proteins were transferred to a nitrocellulose membrane which was then blocked with 1X blocking solution (1X TBS and 5% milk powder) at RT for 1 h. The membrane was then incubated with 1/2,000 biotin-conjugated rabbit anti-HBsAg antibody (Abcam) and 1/100,000 mouse anti-β-tubulin I antibody (T7816; Sigma-Aldrich, Saint Louis, MO, USA), the housekeeping control, at 4°C overnight. The membrane was then washed and stained with 1/20,000 IRDye^®^ 800CW goat anti-rabbit antibody and 1/20,000 IRDye^®^ 680RD goat anti-mouse antibody (LI-COR Biosciences, Lincoln, NE, USA) and the signal was acquired using an Odyssey^®^ DLx instrument (LI-COR). Images were analyzed using Empiria Studio^®^ Software 2.3 (LI-COR).

### Mouse immunization

To evaluate the immune response elucidated by the mRNA-LNP vaccine candidates, C57BL/6Ncr mice were immunized i.m. on days 0 and 28. Whole blood was collected on day 1 prior to the prime and on day 28 prior to the boost. On day 42, the termination of the experiment, blood was collected by intracardiac puncture under anesthesia (120 mg ketamine and 16 mg xylazine per Kg body weight). Following the final blood collection, mice were euthanized by cervical dislocation under anesthesia and the spleens were dissected for functional analysis. Seven groups of mice (n= 8) were immunized with: (i) 5 μg per dose of *HBsAg* mRNA-LNP; (ii) 5 μg per dose of *SP-HBsAg* mRNA-LNP; (iii) PBS control; (iv) 5 μg per dose of *Luc* mRNA-LNP as irrelevant mRNA control; (v) 1 μg per dose of ENGERIX^®^-B vaccine (GSK). The last two animal groups were immunized with a heterologous immunization schedule: (vi) 1 μg per dose of ENGERIX^®^-B vaccine (GSK) as prime and 5 μg per dose of *HBsAg* mRNA-LNP as boost; and (vii) 1 μg per dose of ENGERIX^®^-B vaccine (GSK) as prime and 5 μg per dose of *SP-HBsAg* mRNA-LNP as boost.

To evaluate the efficacy of the vaccines, C57BL/6 mice were first immunized on days 0 and 28 i.m. On day 56 after priming, mice were challenged with 5 x 10^9^ of rAAV8-1.3HBV viral genome/mL. Each animal was injected i.v. with 200 μL (1 x 10^9^ AAV/HBV) via tail vein and the experiment termination was set on day 70. Whole blood was withdrawn on days 63 and 70; and spleens were dissected at termination for further functional analysis. Three groups of mice (n= 8) were immunized with: (i) 5 μg per dose of *HBsAg* mRNA-LNP; (ii) 5 μg per dose of *SP-HBsAg* mRNA-LNP; and (iii) PBS control.

To assess the immunological protection exerted by the mRNA vaccines on a CHB mouse model, HBV-carrier mice were generated by injecting C57BL/6 mice with 5 x 10^9^ rAAV8-1.3HBV viral genomes/mL as indicated for the challenge experiment on day -28 before prime. HBV-carrier animals were immunized on days 0 (prime), 14 (boost 1) and 28 (boost 2) i.m. Experiment termination was set on day 73 and whole blood was withdrawn on days -1, 7, 14, 21, 28, 35, 42, 49, 56, 63 and 73. Spleens were dissected at termination for further functional analysis. Four groups of mice (each n= 6) were immunized with: (i) control, injected with PBS, (ii) 10 μg per dose of *HBsAg* mRNA-LNP, (iii) 10 μg per dose of *SP-HBsAg* mRNA-LNP, and (iv) a combinatorial approach priming with 10 μg *HBsAg* mRNA-LNP, followed by first boost with a mix of 5 μg *HBsAg* mRNA-LNP and 5 μg *SP-HBsAg* mRNA-LNP; and finally a second boost with 10 μg *SP-HBsAg* mRNA-LNP.

### Secondary lymphoid organ cell isolation and stimulation

The spleens were dissected from euthanized mice and disaggregated by passing through a nylon Cell strainer (70 μm). Splenocyte suspensions were washed, and the red blood cells (RBC) were lysed by treating with RBC lysis buffer (ThermoFisher) for 5 min at RT. The RBC-free spleen cells were washed and counted. To evaluate the HBsAg-specific T cell responses elicited by immunization, splenocytes were seeded into 96-well U-bottom culture plates at a density of 5 x 10^5^ cells/100 μL/well in RPMI-1640 medium (Life Technologies Ltd.; Scotland, United Kingdom) supplemented with 10% iFBS, 1% penicillin-streptomycin and 2% L-glutamine. Subsequently, the cells were either left unstimulated or stimulated by the addition of 2 μg/mL/peptide HBsAg peptide pool per well and incubated at 37 °C overnight at 5% CO_2_ for intracellular cytokine staining and for cytokine level measurements in supernates. Concanavalin A (2.5 μg/mL) treated splenocytes served as the positive control.

### Flow cytometric analyses

The hmoDCs, transfected or stimulated for 24 hours, were detached as detailed above, collected and washed with FACS buffer. After blocking the Fc receptors with Privigen® Immunoglobulin solution, the cells were stained at 4°C for 30 min with fluorochrome-conjugated anti-human monoclonal antibodies specific for the following determinants and analyzed by flow cytometry: HLA-DR (APC), CD86 (BD Horizon^TM^ V450), CD80 (PE) and CD40 (PE-Cy^TM^7) (BD Biosciences) (Table S1). Stained cells were washed with FACS buffer and stained with 7-AAD (BD Biosciences) viability dye at RT for 5 min in the dark before acquisition. The hmoDCs stained cells were acquired with an LSR II flow cytometer (BD Biosciences) and analyzed with FlowJo software version 10.8 (Becton, Dickinson and Company).

For experiments involving the activation DCs and T cells *in vivo*, 10^6^ freshly-isolated splenocytes and iLN cells were washed with FACS buffer and the FC receptors were blocked by incubation with anti-CD16/CD32 monoclonal antibody (eBioscience, San Diego, CA, USA). Fluorochrome-conjugated anti-mouse monoclonal antibodies (Table S1) were used to stain the following determinants at 4°C for 30 min in the dark.

Splenocytes stimulated overnight were evaluated by ICS to detect the HBsAg-specific T cell responses. The protein transport inhibitor Brefeldin A was added 15 hours before cell analysis to block secretion and enhance the detection of intracellular cytokines. The cells were collected and washed with FACS buffer; the Fc receptors were blocked by treatment with anti-CD16/CD32 monoclonal antibody (eBioscience) and L/D dye (FVS700) was added for viability staining. Subsequently, cells were stained for 30 min at 4°C with fluorochrome-conjugated anti-mouse monoclonal antibodies specific for the following extracellular determinants: CD8α (BV510) purchased from BD Biosciences, and CD3ε (APC-Cy7) from BioLegend. After fixation and permeabilization with the BD Cytofix/Cytoperm^TM^ Fixation/Permeabilization Kit (BD Biosciences), cells were stained intracellularly for 30 min at 4°C with the following fluorochrome-conjugated anti-mouse monoclonal antibodies: IL-2 (APC) purchased from BioLegend, TNF-α (FITC) from Invitrogen, IFN-γ (BV421), and CD4 (PE-Cy7) from BD Biosciences (Table S1). All stained-splenocytes were acquired with a FACSAria III flow cytometer (BD Biosciences) and analyzed with FlowJo software version 10.8 (Becton, Dickinson and Company).

### Cytokine and chemokine quantitation

Transfected/stimulated hmoDCs culture supernates were collected after 24 hours, and the concentrations of cytokines and chemokines were quantified using the LEGENDplex Human Essential Immune Response Panel (BioLegend) following the manufacturer’s instructions. The panel included the following human cytokines/chemokines: IL-2, CXCL10 (IP-10), IL-1β, TNF-α, CCL2 (MCP-1), IL-17A, IL-6, IL-10, IFN-γ, IL-12p70, IL-8 and free active TGF-β1. Similarly, the supernates were collected from cultures of mouse splenocytes following 24 hours stimulation and the concentration of the following cytokines was determined using the LEGENDplex mouse Th Cytokine Panel (BioLegend): IFN-γ, IL-2, TNF-α, and IL-6. Samples derived from both human and mouse cell cultures were acquired with an LSR II flow cytometer (BD Biosciences) and data were analyzed using the Legendplex Data Analysis Software Suite (BioLegend).

### Serological analyses

The sera were collected from whole blood and anti-HBsAg antibodies were quantified by enzyme linked immunosorbent assay. Anti-mouse IgG H&L (HRP) (ab6789; Abcam) was used to titer total IgG in serial dilutions of the sera using regression analysis. HRP-conjugated goat anti-mouse IgG1 (ab97240; Abcam) and goat anti-mouse IgG2c (HRP) (ab97255) were used to calculate the IgG2c/IgG1 ratio from the optical density obtained at the same fixed dilutions of sera. For the challenge and the CHB mouse model, whole blood samples were collected in K_2_-EDTA coated tubes and plasma samples were obtained after centrifugation at 7,000 x *g*, 4°C for 10 min to quantify plasma anti-HBsAg antibodies, HBsAg and HBeAg. Plasma anti-HBsAg antibodies were measured using a chemiluminescent microparticle immunoassay (CLIA, Autobio, 30040303CM01). Plasma HBsAg was quantified using the HBsAg Quantitative CLIA (Autobio, CL 0310). Mouse plasma samples were diluted 20-fold and HBsAg was quantified according to the manufacturer’s instructions. Plasma HBeAg was quantified after 6-fold dilution of the plasma using the HBeAg CLIA kit (Autobio, CL 0312) following manufacturer’s instructions.

### Statistical analyses

All graphs were created, and data were analyzed using GraphPad Prism version 9.4.1 (GraphPad Software Inc., San Diego, CA, USA). A two-tailed Student’s *t*-test was used to compare two groups; a one-way ANOVA test followed by Tukey’s multiple comparison test and a two-way ANOVA test followed by Tukey’s multiple comparison test were used to compare three or more groups. All values are expressed as mean ± SEM; statistically significant *p*-values were: *, p <0.05; **, p <0.01; ***, *p* <0.001; ****, *p* <0.0001.

## Acknowledgements

The authors are grateful to Stephen H. Gregory (Providence, RI, USA) for editing this manuscript.

## Authors’ Contributions

M.L.C. and S.G. conceptualized the study. N.P. provided oversight on the study design and provided mRNA vaccines. M.L.C., S.G., N.P. and M.J.L. designed vaccines. M.L.C., S.G., and M.J.L. designed and planned experiments. M.J.L. performed *in vitro* experiments. M.L.C., M.J.L., G.A.I., and I.R.B. formulated mRNA into LNPs. M.L.C., M.J.L., M.S., R.G., and L.P. performed *in vivo* experiments. D.F. assisted with in vivo experiments. M.J.L., M.S., R.G., and L.P. processed animal samples and performed flow cytometry analyses. M.J.L., R.G., and S.F. contributed to ELISA experiments. M.B. assisted with experimental design. M.L.C. and M.J.L. analyzed data and performed statistical analyses. M.L.C. prepared figures and the manuscript. S.G. and N.P. edited the manuscript. S.G. supervised the study and acquired funding. All authors performed critical revisions and approved the final manuscript text and figures.

## Competing Interests

M.L.C. is currently an employee at BioNTech SE (Mainz, Germany), however, the contributions from M.L.C. were made prior to his employment at BioNTech SE. The company had no role in the design of the study; in the collection, analysis, or interpretation of data; in the writing of the manuscript; or in the decision to publish the results. N.P., M.L.C., M.J.L. and S.G. have a provisional patent application that describes the use of nucleoside-modified mRNA in lipid nanoparticles to treat and/or prevent HBV infections. The remaining authors declare no conflict of interest.

