## Supplementary Material for "mRNA-LNP vaccines against Hepatitis B virus induce protective immune responses in preventive and chronic mouse challenge models"

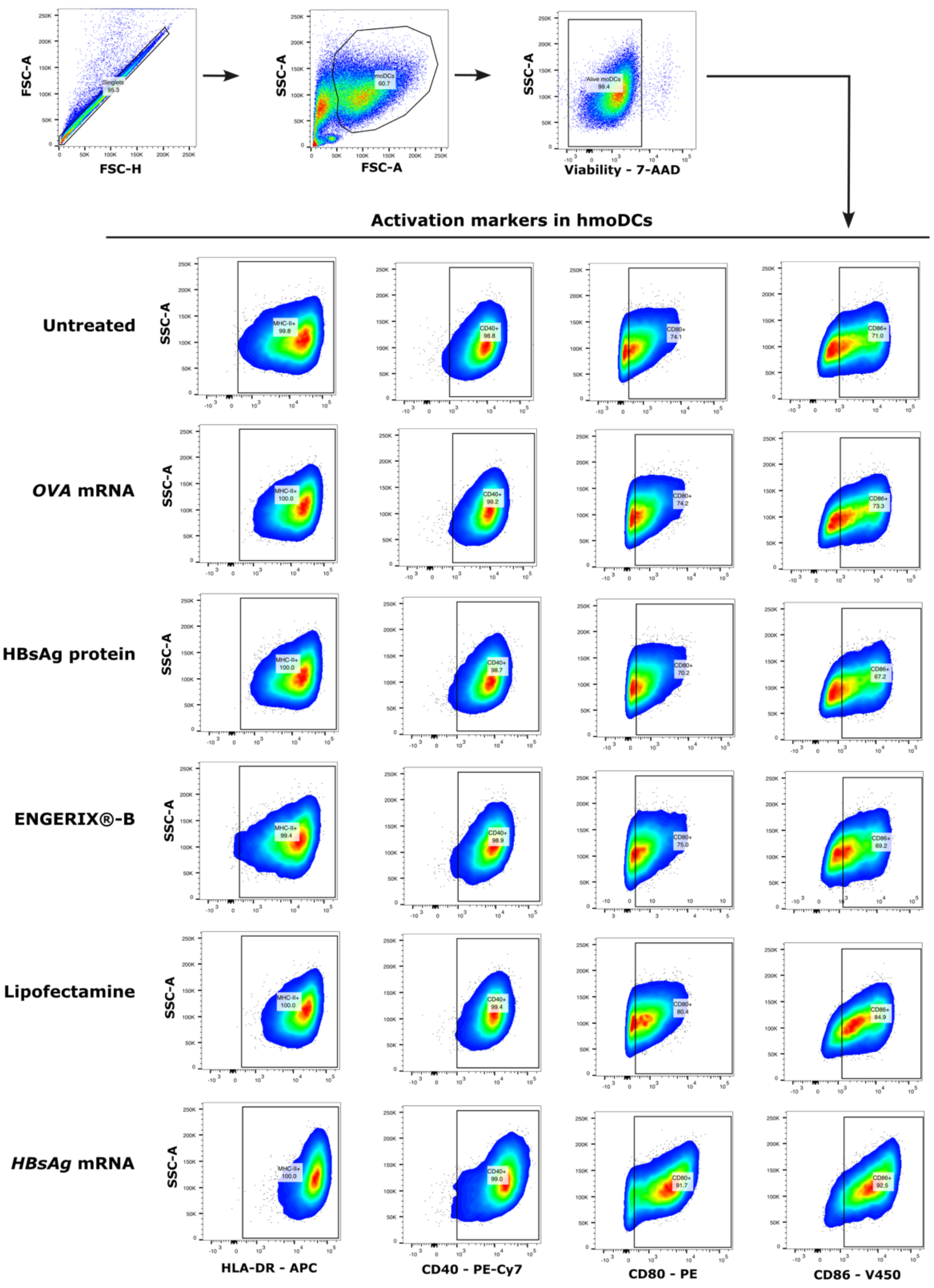


**Figure S1. Gating strategy of the hmoDC transfection.** Flow cytometry dot plots of the unstimulated and transfected hmoDCs with *HBsAg* mRNA to study their activation through HLA-DR, CD40, CD80 and CD86 surface markers. OVA mRNA, HBsAg protein, ENGERIX®-B and Lipofectamine served as controls.

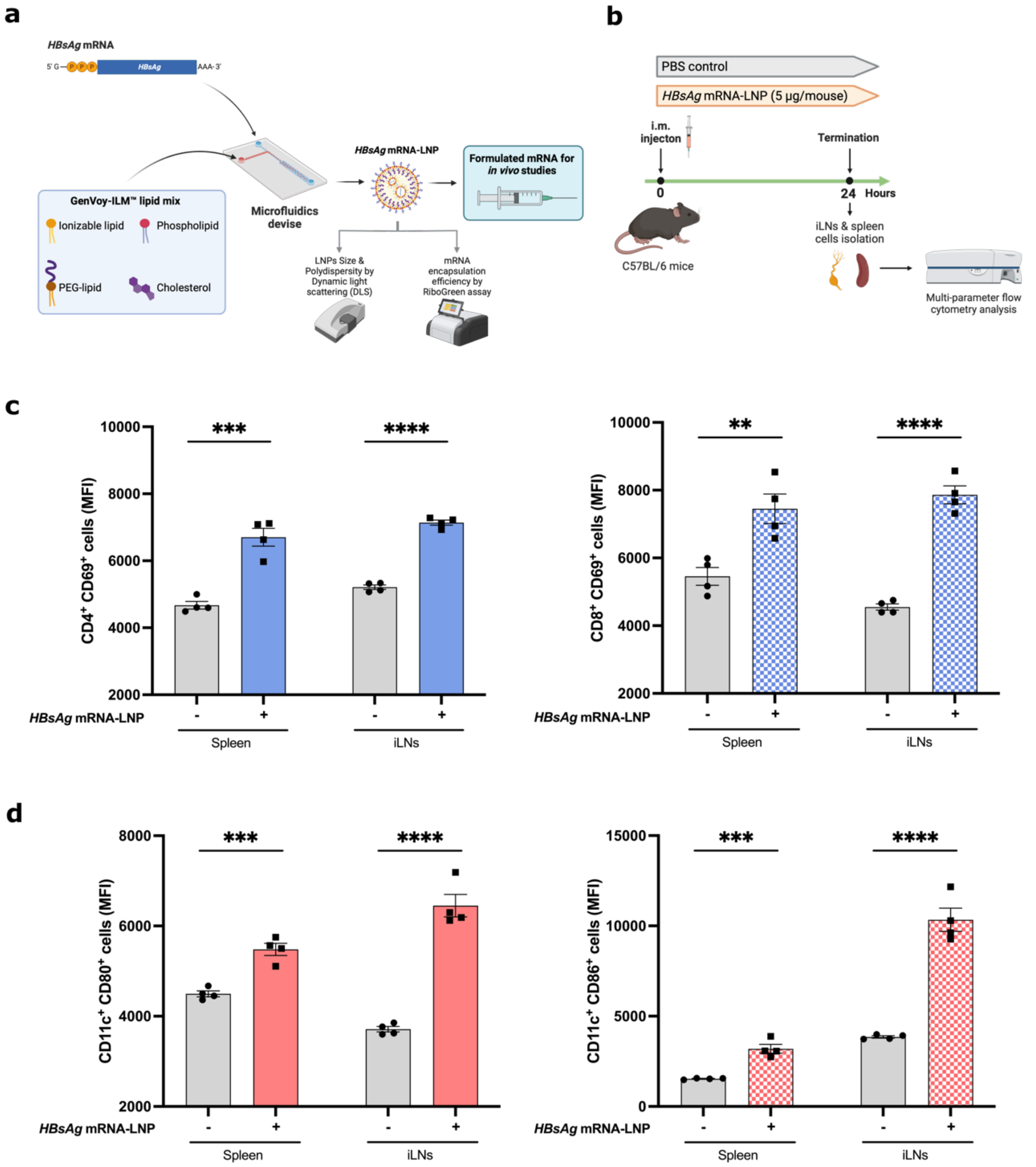


**Figure S2: *HBsAg* mRNA formulated in LNP induces early activation of DCs and T cells in secondary lymphoid organs.** (a) Schematic representation of mRNA-based vaccine preparation. *HBsAg* mRNA was formulated in LNP, and mRNA-LNP were then characterized. (b) Experimental design. Mice were vaccinated i.m. with 5 µg *HBsAg* mRNA-LNP. Spleen and inguinal lymph nodes were dissected for evaluation 24 hours after injection. Expression of early activation markers (c) CD69 by $CD4^+$ and $CD8^+$ T cells, and (d) CD80 and CD86 by DCs isolated from the spleen and lymph nodes were compared to the expression of cells derived from the PBS control group. Data are the means ± SEM for animal cohorts

(4 mice/group). Significantly different from the control group: **p< 0.01, ***p< 0.001; ****p <0.0001 (two-tailed Student's *t*-test).

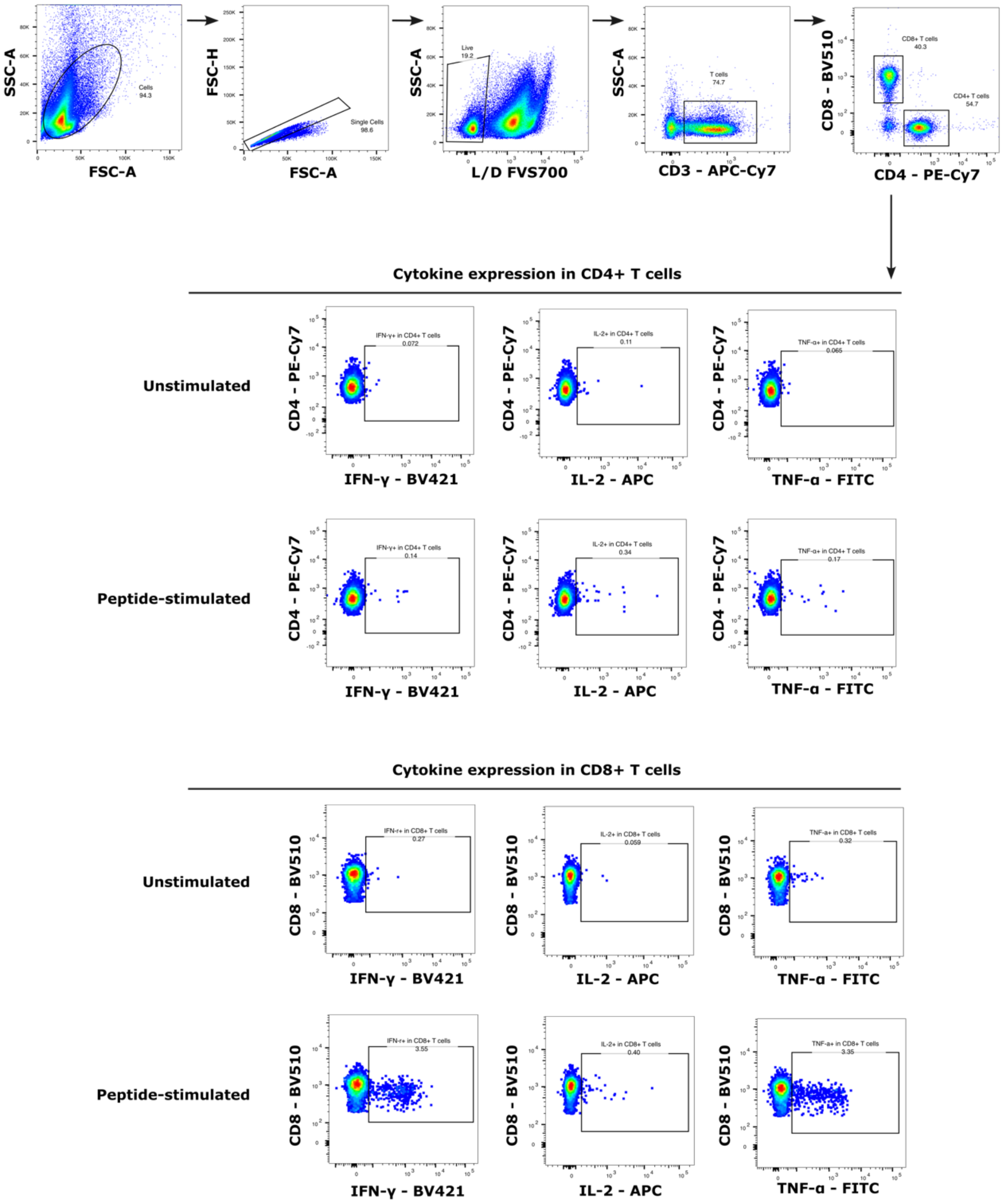


**Figure S3. Gating strategy of the intracellular cytokine staining.** Flow cytometry dot plots of the unstimulated and peptide-stimulated splenocytes to study the specific CD4$^{+}$ and CD8$^{+}$ T cell responses in immunized mice.

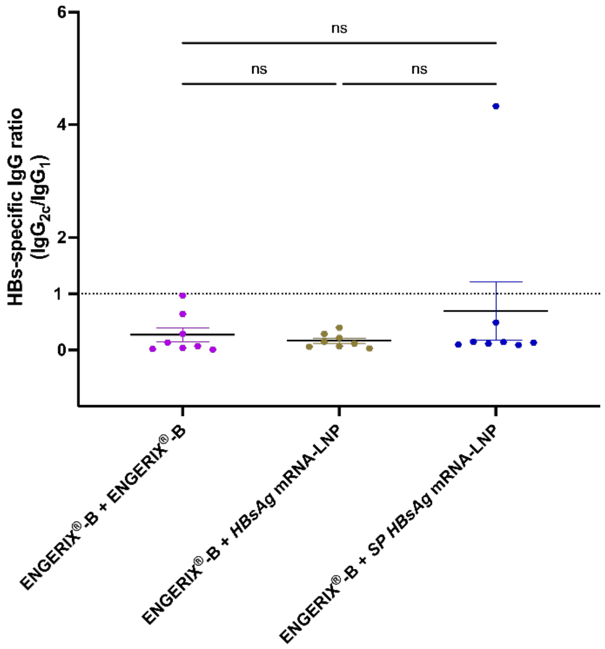


**Figure S4: Anti-HBsAg IgG subtypes antibodies from mice immunized with the heterologous immunization schedule.** IgG2c/IgG1 ratio at termination (d42). Data are the means ± SEM for animal cohorts (8 mice/group). Significance tested compared to the other groups: no significant differences (ns) (one-way ANOVA, Tukey's multiple comparisons test).

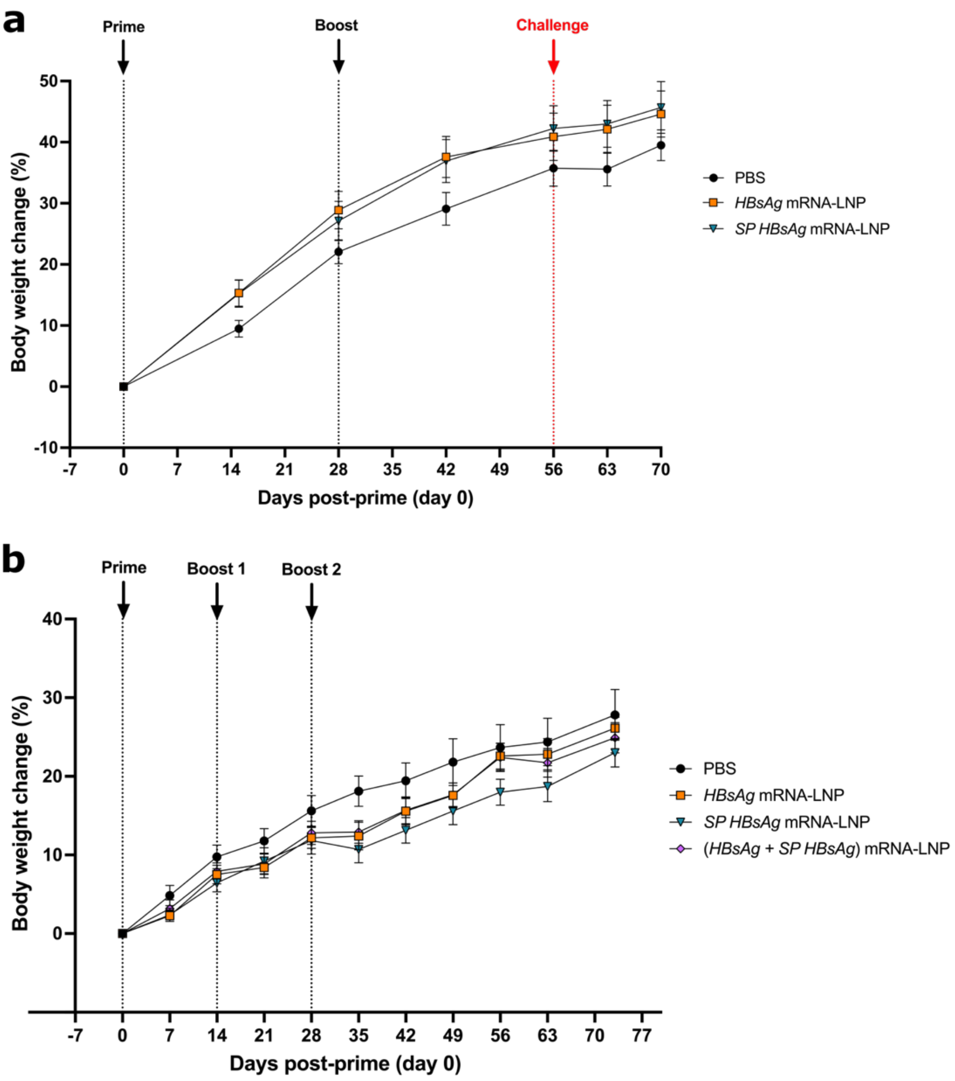


**Figure S5. Body weight change in immunized mice.** Evaluation of the reactogenicity by measuring the percentage of the body weight change in rAAV8-1.3HBV challenged mice (a) and in CHB mice (b).

**Table S1: List of antibodies used for flow cytometry analysis.**

| Antibody | Fluorochrome | Reference | Clone | Dilution |
|---|---|---|---|---|
| **Activation markers of transfected/stimulated hmoDCs** | | | | |
| HLA-DR | APC | 559866 | G46-6 | 1/10 |
| CD86 | BD Horizon$^{TM}$ V450 | 560357 | 2331 (FUN-1) | 1/25 |
| CD80 | PE | 557227 | L307.4 | 1/10 |
| CD40 | PE-Cy$^{TM}$7 | 561215 | 5C3 | 1/25 |
| ***In vivo* DC and T cell activation** | | | | |
| CD8a | Super Bright$^{TM}$ 436 | 62-0081-82 | 53-6.7 | 1/125 |
| CD3 | eFluor$^{TM}$ 506 | 69-0032-82 | 17A2 | 1/60 |
| CD80 | PerCP-eFluor$^{TM}$ 710 | 46-0801-82 | 16-10A1 | 1/200 |
| CD4 | FITC | 553047 | L3T4 | 1/800 |
| CD69 | PE-Cy$^{TM}$7 | 552879 | H1.2F3 | 1/80 |
| CD11c | PE | 117308 | N418 | 1/100 |
| CD86 | APC | 105012 | GL-1 | 1/100 |
| **Intracellular cytokine staining** | | | | |
| IL-2 | APC | 503810 | JES6-5H4 | 1/80 |
| IFN-γ | BD Horizon$^{TM}$ BV421 | 563376 | XMG1.2 | 1/100 |
| TNF-α | FITC | 11-7321-82 | MP6-XT22 | 1/200 |
| CD8α | BD Horizon$^{TM}$ BV510 | 563068 | 53-6.7 | 1/200 |
| CD3ε | APC-Cy$^{TM}$7 | 100330 | 145-2C11 | 1/40 |
| CD4 | PE-Cy$^{TM}$7 | 563933 | GK1.5 | 1/200 |
